# An annotated reference library for supporting DNA metabarcoding analysis of aquatic macroinvertebrates in French freshwater environments

**DOI:** 10.1101/2024.09.23.614429

**Authors:** Paula Gauvin, David Eme, Isabelle Domaizon, Frédéric Rimet

## Abstract

Freshwater ecosystems are increasingly threatened by human activities, leading to biodiversity loss and ecosystem degradation. Effective biodiversity monitoring, particularly through the use of aquatic macroinvertebrates as bioindicators, is crucial for assessing ecological health. While traditional morphological methods face limitations, DNA metabarcoding offers higher accuracy and efficiency in species identification using environmental DNA. However, the success of metabarcoding is contingent on the quality of reference libraries, which are often incomplete or biased. This study aimed to construct a comprehensive COI-based DNA barcode library for freshwater macroinvertebrates in France, specifically targeting short gene regions amplified with fwhF2/fwhR2N primers, suitable for degraded DNA. A list of species occurring in French freshwater ecosystems was established from official national checklists and Alpine lake surveys. The resulting library was analyzed for taxonomic completeness, barcode coverage and genetic diversity. The checklist consisted of 2,841 species across 10 phyla, for which 56% had at least one COI-5P sequences available in the Barcode of Life Data System (BOLD). Alignment challenges with the primers were identified for certain taxa, particularly among Coleoptera, Diptera, and Malacostraca. The genetic diversity approached by the number of haplotypes per species highlighted that most of the species have limited diversity, with only 3 species having more than 100 haplotypes. Finally, this study showed that a total of 57 haplotypes were shared among 116 distinct species. This work emphasized the need for expanded sequencing efforts to improve barcode coverage and highlighted the pitfalls associated with the use of these primers for further biodiversity assessment of macroinvertebrates with degraded DNA.

## Introduction

Freshwater ecosystems face escalating anthropogenic pressures threatening their functionality (Reid et al. 2019) and leading to significant biodiversity loss (Young et al. 2016, Borgwardt et al. 2019). In this context, biodiversity monitoring is crucial to provide relevant ecological diagnoses and support the management and conservation of these ecosystems and the vital services they provide. Aquatic macroinvertebrates are widely used as bioindicators due to their abundant presence, high species diversity and sensitivity to both anthropogenic and natural disturbances (Hering et al. 2006). These organisms primarily inhabit the littoral zone and play pivotal roles in community assembly and food web dynamics, contributing significantly to beta diversity and metacommunity structure across diverse aquatic habitats (May 2019). Consequently, they serve as key indicators in environmental monitoring programs, facilitating assessments of the ecological status of aquatic ecosystems as mandated by the EU Water Framework Directive (WFD) and national legislation (Mondy et al. 2012). The standardized methodology for macroinvertebrate biomonitoring in Europe is based on morphological identification through binocular microscopy (ANFOR, 2010; CEN 2019). This methodology encounters significant challenges, including the labor-intensive and expensive nature of collecting and sorting individual benthic invertebrates, which hinders its broader adoption in routine biomonitoring (Blackman et al. 2019; Bonada et al. 2006). Moreover, routine morphological identification poses drawbacks such as high expertise demands and lower taxonomic resolution (Leese et al. 2016, 2018, Bean et al. 2017, Hering et al. 2018). Notably, many taxa can only be identified at limited taxonomic resolution (Caesar et al. 2006), without a timely effort of highly skilled taxonomists. This is the case for Chironomids, a common group in freshwaters, frequently classified at the family level due to challenges in species or genus identification (Beermann et al. 2018). Additionally, immature life stages lacking diagnostic morphological traits hinder accurate species identification and therefore limit the proper quality assessment of freshwater ecosystems (Sweeney et al. 2011). These challenges underscore the need for innovative approaches to enhance the efficiency and accuracy of macroinvertebrate biomonitoring practices. The recent advent of DNA metabarcoding techniques, integrating amplicon barcoding with high-throughput sequencing, represents a promising advancement in biomonitoring applications (Deiner et al. 2017, Pawlowski et al. 2018, Carraro and Altermatt 2024), providing a valuable complement to morphology-based approaches for species identifications. Metabarcoding finds application in analyzing environmental DNA (eDNA) samples obtained from water, biofilm or sediment (Sakata et al. 2020; Valentini et al. 2016, Rivera et al. 2021, Gauvin et al. submitted) or DNA extracts derived from a homogenate of the sample’s fauna (Taberlet et al. 2012). These molecular innovations enable simultaneous processing of multiple samples, identification of small taxa, immature or larval stages, and offer increased sensitivity and specificity, often revealing hidden diversity, while enhancing time and cost effectiveness (Shokralla et al. 2012, Pochon et al. 2015, Holman et al. 2019). These advantages facilitate direct comparison among sites and studies and enable higher spatial-temporal frequency in monitoring due to increased throughput (Bush et al. 2019). The efficiency of macroinvertebrates DNA metabarcoding relies heavily on primer sets’ effectiveness across a broad taxonomic spectrum. The recovery rate of taxa using metabarcoding is contingent upon various factors, including the taxonomic resolution of the gene marker employed (e.g. COI or ribosomal markers like 16S, e.g, Elbrecht et al. 2016), amplicon length (Meusnier et al. 2008), primer universality, and the number of primer pairs utilized to amplify target taxonomic groups (Elbrecht and Leese 2015, Gibson et al. 2015). Specifically, a segment of the cytochrome c oxidase subunit I (COI) gene has emerged as the standard DNA barcoding marker for most animal groups (Hebert et al. 2003a), with over 95% of species in diverse animal groups exhibiting distinctive COI sequences in test assemblages (Hebert et al. 2003b, 2004). Given the considerable phylogenetic diversity among macroinvertebrates, the high taxonomic resolution and existing reference data for the COI marker, make it a judicious choice for metabarcoding freshwater macroinvertebrate communities (Ratnasingham and Hebert 2007, Andújar et al. 2018). Furthermore, well-established barcoding gaps for the COI marker in freshwater macroinvertebrates underscore its suitability for such applications (Zhou et al. 2009). Numerous primers of COI gene of varying base pair sizes are available in the literature for DNA amplification of macroinvertebrates communities. Environmental DNA released by target taxa can degrade rapidly (Seymour et al. 2018). Consequently, for amplifying degraded DNA, that is extracted from water samples for example, targeting a short COI marker region is suggested to enhance amplification success (Barnes et al. 2014, Jelger Herder et al. 2014). The effectiveness of DNA metabarcoding programs relies on open, comprehensive and accurate reference sequence libraries (Briski et al. 2016, Oliveira et al. 2016, Weigand et al. 2019), ensuring precise taxonomic assignment (Richardson et al. 2018, Rodriguez‐Ezpeleta et al. 2021). In that sense, incomplete DNA barcode libraries, as in poorly represented species, pose a significant challenge, often leading to false negatives and compromising biodiversity assessments (Ardura 2019, Leite et al. 2020, Duarte et al. 2021). Therefore, evaluating these gaps and the quality of sequence data in reference libraries is imperative for the effective implementation of DNA-based tools in biodiversity assessments (Duarte et al. 2020). Given the significance of reference libraries completeness, primarily those that are pertinent in the context of the WFD, numerous studies have been conducted to assess their representativeness by comparing them with lists of described species (Trebitz et al. 2015, Weigand et al. 2019, Duarte et al. 2020, Leite et al. 2020, Specchia et al. 2020, Csabai et al. 2023). Despite efforts to assess their representativeness, biases in taxonomic coverage persist within reference libraries (Li et al. 2019, Weigand et al. 2019). Indeed, numerous studies have emphasized the construction of tailored reference libraries to suit the geographic context of the research area (Ficetola et al. 2021, Mugnai et al. 2023) and emphasized the importance of such databases to be refined with custom sequences specific to local study areas and free from unexpected taxa. Additionally, research by Abad et al. 2016 and Schenekar et al. 2020 has highlighted the value of possessing DNA barcodes for local species. These studies have demonstrated that DNA barcodes from local organisms can improve the accuracy of taxonomic assignments and reveal previously unrecognized biodiversity, leading to adjustments in taxonomic classifications among species. Our study is in line with the recommendations proposed by Blackman et al. 2023, which advocate for compiling a comprehensive list of target species within the study area and assembling accurate sequences pertinent to those species. Our first objective is to assemble a list of macroinvertebrate species known to be present in French freshwater ecosystems. The second objective is to construct a DNA barcode library utilizing COI primers, particularly focusing on a short region of this gene which is highly effective when targeting degraded DNA (fwhF2/fwhR2N, 205pb, Vamos et al. 2017), to identify freshwater macroinvertebrate fauna in France using metabarcoding techniques and to assess its genetic completeness. Our third objective is to identify the gaps of this new reference library, thereby providing insight into its limitations and facilitating data interpretation for futures users.

## Material and methods

### Checklist constitution for French metropolitan freshwater macroinvertebrates species

In order to establish a checklist of macroinvertebrates taxa of freshwater aquatic habitats in metropolitan France, three checklists were complied. The first one comes from PERLA, an interactive tool managed by a French regional environmental agency (DREAL Auvergne-Rhône-Alpes) accessible at http://www.perla.developpement-durable.gouv.fr/index.php. PERLA serves as a national comprehensive checklist for water managers and encompasses larvae, nymphs and adults across various taxonomic groups, including insects, molluscs, crustaceans, and worms, among others, found in rivers and aquatic ecosystems. In the manuscript, we will refer to this checklist as the ‘French aquatic ecosystems checklist’. The second checklist is coming from macroinvertebrate surveys conducted in four Alpine lakes (Lake Geneva, Lake Annecy, Lake Bourget and Lake Tignes) from 2015 to 2022. In the manuscript, we will refer to this checklist as the ‘French Alpine lakes checklist’. The third checklist we used was from a French NGO, Opie-benthos (https://www.opie-benthos.fr/opie/monde-des-insectes.html), studying freshwater insect’s taxonomy and diversity. Currently, Opie-benthos leads comprehensive inventories across France, specifically targeting insect species undergoing a part of their life cycle within diverse aquatic ecosystems. In the manuscript, we will refer to this checklist as the ‘French aquatic insect’s species list’. These three checklists were merged into a single one. When the taxonomic resolution of taxa was limited to a level above species (genus, family, class, phylum), we consulted the taxonomic referential of the National Inventory of Natural Heritage (TAXREF v17.0 2024, available at https://inpn.mnhn.fr/telechargement/referentielEspece/taxref/17.0/menu) to select species from those ranks. The phylum Acanthocephala and Rotifera, the classes Copepoda and Ostracoda (Arthropoda, Crustacea) and the families Succineidae (Arthropoda, Insecta, Diptera) and Hydrachnidae (Arthropoda, Arachnida, Trombidiformes), initially listed in French aquatic ecosystems checklist, were omitted from our search from TAXREF v17.0 2024 as they are not considered as freshwater macroinvertebrates according Tachet et al. 2000. Once those macroinvertebrates species were retrieved for each taxonomic levels without species identification, several filters from TAXREF were applied to select only French metropolitan freshwater species from this taxonomic referential. First, we selected only the species occurring in the following habitats: freshwater, marine and freshwater, brackish water, continental (land and freshwater). Then, from this updated list, only species occurring in metropolitan France were kept. Finally, species characterized by one those 8 status over 15 status in total were selected (present (native or undetermined), endemic, sub-endemic, cryptogenic, introduced, invasive introduced, non-established introduced (including cultivated / domestic), occasional). The checklists described above and the summarized species list resulting from these compilations, inclusive of all species from the mentioned inventories, is referred as the “French metropolitan freshwater macroinvertebrates species list” and all together are available at https://doi.org/10.57745/LMOXEW in the ‘Checklists composition data’ file for future reference.

### Sequences origin and cleaning steps

All the sequences for those species listed in the French metropolitan freshwater macroinvertebrates species list, whatever the gene, were downloaded from October 2023 to May 2024, from the Barcode of Life Data System v4 -Bold-repository (https://boldsystems.org/index.php/databases). We opted to utilize Bold as reference library instead of GenBank due to concerns regarding the latter’s unverified submission process, which frequently results in misannotated sequences (Kozlov et al. 2016, Locatelli et al. 2020, Steinegger and Salzberg 2020). The presence of stop codons were checked among the retrieved sequences and none were found. The absence of sequences for a given species may indicate either that the search in BOLD returned “Unmatched terms” or that while the species was found, no sequences were publicly available, rendering them private sequences (regardless of the reason for their absence, they were all defined as “not available sequences” for this analysis). Before concluding that a species lacked records in BOLD for the species from the French aquatic ecosystems checklist and French Alpine lakes checklist, synonyms were searched using INPN. The details of taxa resulting from this taxonomic harmonization are available in Table S1. From this file containing all the sequences retrieved (whatever the gene) for the queried species of the French metropolitan freshwater macroinvertebrates species list, the COI-5P sequences belonging to the COI gene were selected. All the remaining sequences were aligned with the fwhF2/fwhR2N primer pair using MAFFT version 7 (Auto) (https://mafft.cbrc.jp/alignment/server/), by genus or species group (as in all sequences assigned to the same genus or specie were aligned together). Then, sequences characterized by gaps, insufficiencies in length relative to the COI barcode standard (shorter than 205 pb), or identical (same genetic sequence for one species) from the sequences file, were removed using Jalview (https://www.jalview.org/). Due to the possibility of identical sequences being shared among different species, certain species may have been excluded during this stage, as a result of sequence overlap with other taxa. To address this issue, the list of species obtained from this step was cross-referenced with the initial species list following the alignment process. This comparison identified any missing species, whose sequences were then realigned individually to ensure their accurate representation. This final reference library containing only genetic sequences capable of being aligned by the fwhF2/fwhR2N primer pair is named ‘Aligned DNA library’ hereafter.

### Graphical analysis

We firstly analysed the taxonomic composition of the French metropolitan freshwater macroinvertebrates species list. Secondly, to highlight any gaps in the availability of sequences in our list of species, the taxonomic coverage of barcodes at various taxonomic levels was assessed. Thirdly, we highlighted the taxonomic composition of species with COI-5P sequences that did not align with fwhF2/fwhR2N primers to reveal which species and taxonomic groups could be misrepresented in metabarcoding studies using those primer pair. Then, to explore the species haplotype diversity, the number of different haplotypes available relative to the number of species by phylum was examined. The categorization of unique barcodes ranging from limited (<5 barcodes) to moderate (5-25 barcodes) to good (>25 barcodes), was adopted from Trebitz et al. 2015, and the species with the highest number of unique sequences (haplotype) exceeding 100 were identified. Finally, in order to demonstrate the ability of the primers used in the aligned DNA library to discriminate each species by a unique barcode, groups of taxa sharing the same haplotype were identified.

## Results and discussion

### Taxonomic composition of the French metropolitan freshwater macroinvertebrates species list and barcoding coverage

The French metropolitan freshwater macroinvertebrates species list is composed by 10 phyla, 16 classes, 50 orders, 222 families, 670 genera, and 2841 species (Table 1).

**Table 1.** Summary of taxonomic composition for each phylum of the the French metropolitan freshwater macroinvertebrates species list.

| Phylum | Number of |  |  |  |  |
| --- | --- | --- | --- | --- | --- |
|  | Class | Order | Family | Genus | Species |
| Annelida | 1 | 2 | 6 | 18 | 40 |
| Arthropoda | 4 | 20 | 135 | 508 | 2469 |
| Bryozoa | 2 | 2 | 6 | 7 | 11 |
| Cnidaria | 1 | 2 | 3 | 3 | 7 |
| Entoprocta | 1 | 1 | 1 | 1 | 1 |
| Mollusca | 2 | 11 | 24 | 52 | 195 |
| Nematoda | 2 | 9 | 42 | 68 | 99 |
| Nemertea | 1 | 1 | 1 | 1 | 1 |
| Platyhelminthes | 1 | 1 | 3 | 7 | 12 |
| Porifera | 1 | 1 | 1 | 5 | 6 |

Figure 1 provides insights into the taxonomic composition, highlighting that the highest diversity is observed within the phylum Arthropoda, followed by Mollusca and Nematoda, with 2469, 195, and 99 species, respectively. On the other hand, taxa such as Entoprocta and Nemerta are represented by only one species (Figure 1, Table 1).

**Figure 1.**
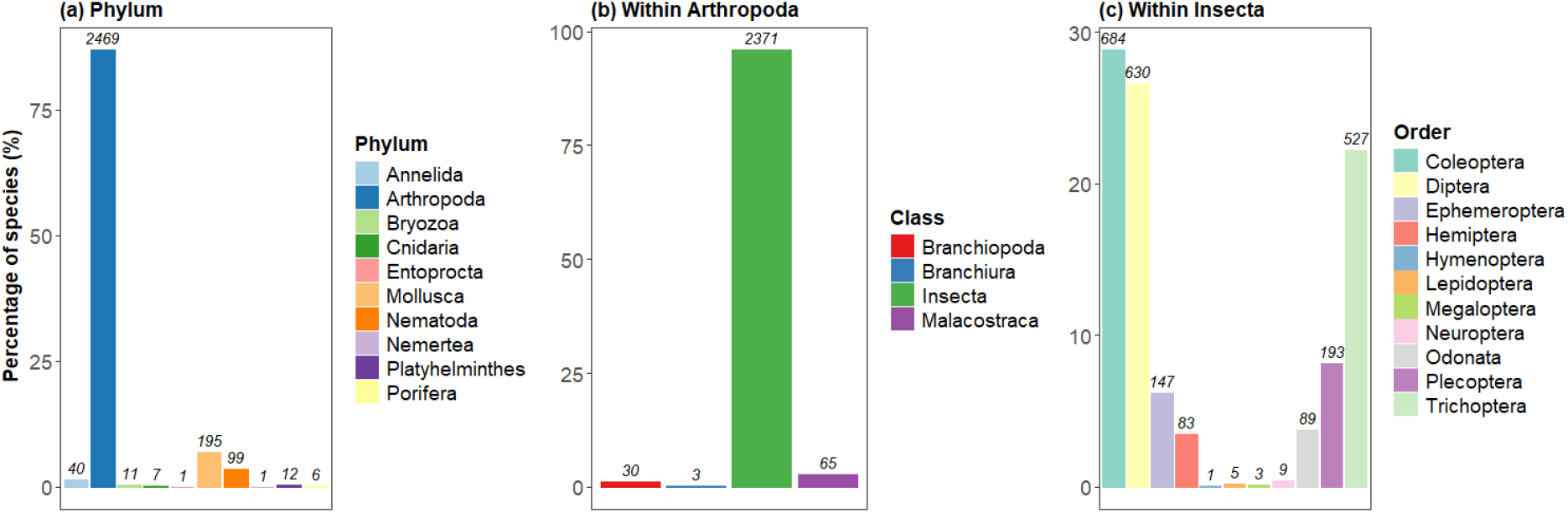
Taxonomic composition of the French metropolitan freshwater macroinvertebrates species list. Percentage of the number of species according to phylum (a), class within the arthropod phylum (b) and order within the insect class (c). The number above each bar represents the total number of species affiliated to each taxonomic rank.

Among the insects, Coleoptera dominate, followed by Diptera and Trichoptera, with 684, 630, and 527 species, respectively. The predominant phyla with the highest number of species in our checklist matches with those identified in a study by Specchia et al. 2020, who conducted a gap analysis of DNA barcodes available in international repositories using the aquatic macroinvertebrate species checklist of a south-eastern Apulia region in Italy. Furthermore, our findings align with the prevailing understanding that Arthropoda, and thus insects, represent the most diverse group of animals, exerting dominance in freshwater ecosystems inventories (Choudhary & Ahi, 2015; Dijkstra et al. 2014; Grosberg et al. 2012; Yeates & Wiegmann, 1999). Moreover, Diptera emerge as the most species-rich group utilized in biomonitoring across Europe, with chironomids recognized for their prevalence and diversity in freshwater habitats (Pinder 1986). Alongside Diptera, Coleoptera (beetles) stand out for their exceptionally high species numbers across various ecoregions and countries in Europe and serving as the most abundant group of aquatic insects (Jäch and Balke 2008, Short 2018). Following these orders, Trichoptera (caddisflies), Plecoptera (stoneflies) and Ephemeroptera (mayflies), collectively known as EPT, emerge as the next three orders in terms of species richness from our checklist. These organisms spend their immature stages in freshwater and are widely employed as biological indicators for freshwater quality assessment (Hering et al. 2004, Sweeney et al. 2011) and ecological investigations, demonstrating robust responses to pollution or climate change (Álvarez-Troncoso et al. 2015). Additionally, Nematoda emerges as a highly diverse group, ranking third in species richness among all phyla listed in our checklist. This outcome may be attributed to the the interest in this phylum is its utility in ecological assessments, as nematodes are used for this purpose since a long time (e.g. Bongers, 1990; Moreno et al. 2011). From all the sequences retrieved based on the species of the French metropolitan freshwater macroinvertebrates species list, 85.4% were belonging to COI-5P gene (Figure S1). This result was expected as BOLD is the main repository for COI sequences (Ratnasingham and Hebert 2007). Overall, 56% of the 2841 species listed in the French metropolitan freshwater macroinvertebrates species list possessed at least one COI-5P genetic sequences publicly available in BOLD database (Figure 2).

**Figure 2.**
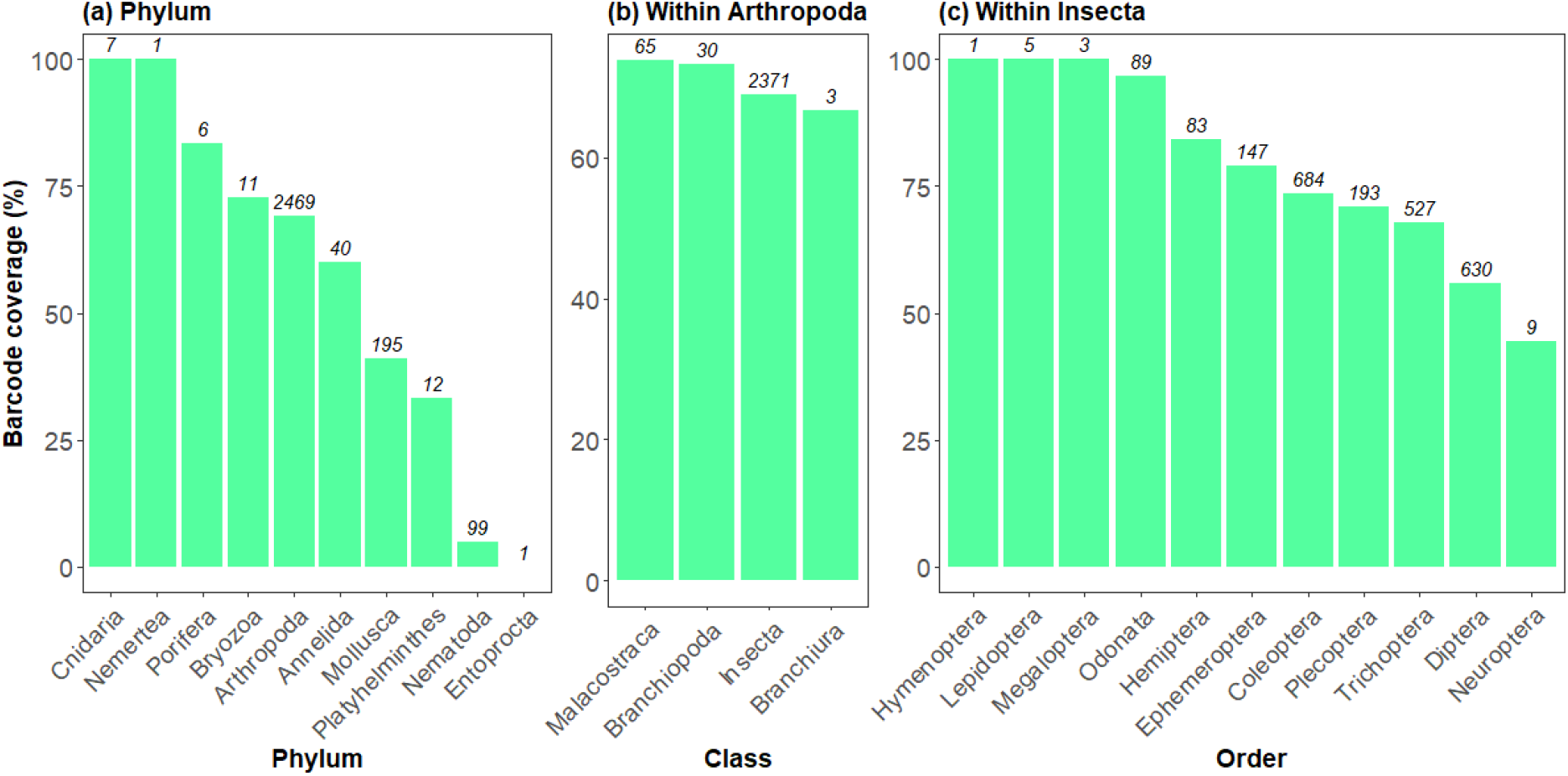
Barcoding coverage of the French metropolitan freshwater macroinvertebrates species list. The barcoding coverage gives for each phylum, for each class within Arthropoda and for each order within Insecta the total percentage of species with available COI-5P public sequence in BOLD. The number above each bar represents the total number of species listed in the Freshwater macroinvertebrate checklist.

There are some exceptions with Nemerta, which have a barcode coverage of 100% and the Entoprocta which have 0% coverage (both phylum with only species referenced). Mollusca (195 species) and Nematoda (99 species), which are the second and third most diverse phylum in terms of species in the French metropolitan freshwater macroinvertebrates species list, have one the worst barcode coverage of all phyla (41% and 5.1%, respectively). Within Arthropoda, most of the class have a barcode coverage above 60%, with the highest being the Malacostraca (65 species) (73.8%) and the lowest the Branchiura (3 species) (66.7%). Finally, within insects, Hymenoptera (1 specie), Lepidoptera (5 species), Megaloptera (3 species), Odonata (89 species) and Hemiptera (83 species) have the highest barcode coverage (>80%). The three most species diverse insects orders (Coleoptera (684 species), Diptera (630 species), Trichoptera (527 species)) have a barcode coverage of 73.4%, 55.9% and 67.9%, respectively (Figure 2). Compared to previous gap analyses carried out in other countries, our analysis in freshwater ecosystems in France revealed a slightly lower barcoding coverage for freshwater macroinvertebrates (56%). For instance, investigations in specific regions such as the Apulia Region of Southeast Italy reported DNA barcode availability for 58% of listed aquatic Macroinvertebrate species (Specchia et al. 2020), while a study in Atlantic Iberia documented coverage for 63% of macroinvertebrates (Leite et al. 2020). Similarly, a comprehensive assessment of 4502 freshwater invertebrate species utilized in ecological quality assessments indicated that 60% possessed one or more barcodes (Weigand et al. 2019). Our findings of the lowest barcode coverage at the phylum level align with previous studies, which also reported very low barcode coverage for freshwater Platyhelminthes, with only three species having sequences deposited in examined databases (two in our study). The low taxonomic coverage for Mollusca in our study can be attributed to the high number of DNA barcodes deposited in GenBank rather than BOLD. As our study exclusively utilized BOLD for sequence retrieval, this likely accounts for the observed low barcode coverage of Mollusca, despite it being the second most listed phylum in our checklist. Additionally, although nematodes are taxonomically diverse and ecologically significant, they are generally overlooked in existing surveys (Weigand et al. 2019 and references therein). Insects exhibited the highest number of available COI sequences, consistent with previous studies highlighting the overrepresentation of Arthropoda, particularly insects (Meglécz, 2023). The barcode coverage for Insecta was 69%, aligning with other findings such as Weigand et al. (2019), who reported that 66% of monitored insect species were barcoded, and Trebitz et al. (2015), who found approximately 70% representation in BOLD for Great Lakes fauna. Within Insecta, Diptera had the lowest coverage on BOLD at 55.9%, while Odonata and Hemiptera were the best covered, with over 80% of species barcoded in each group, similar to Weigand et al. (2019). The large number of DNA barcodes accessible within public databases often reflects the intensity of dedicated studies and associated barcoding projects. Certain taxonomic groups, such as Arthropoda or Mollusca receive disproportionate attention, resulting in a heightened focus and increased deposition of sequences within genetic databases (Briski et al. 2011, 2016, Ardura 2019). Consequently, the absence of comprehensive reference databases may lead to false-negative outcomes (Klymus et al. 2017), while inaccuracies within these databases can engender false-positive identifications, as evidenced by instances of misreporting species presence (Port et al. 2016). Furthermore, the false assignment of sequences to closely related species may ensue when references for the true species are absent (Schenekar et al. 2020, Couton et al. 2022). Resolving this issue necessitates to sequence new specimens of target taxa and their subsequent integration into reference libraries.

### Gap analysis of the aligned DNA library

The Figure 3 represents the taxonomic composition of the 27 species from the French metropolitan freshwater macroinvertebrates species list which cannot be aligned with the primer pair fwhF2/fwhR2N.

**Figure 3.**
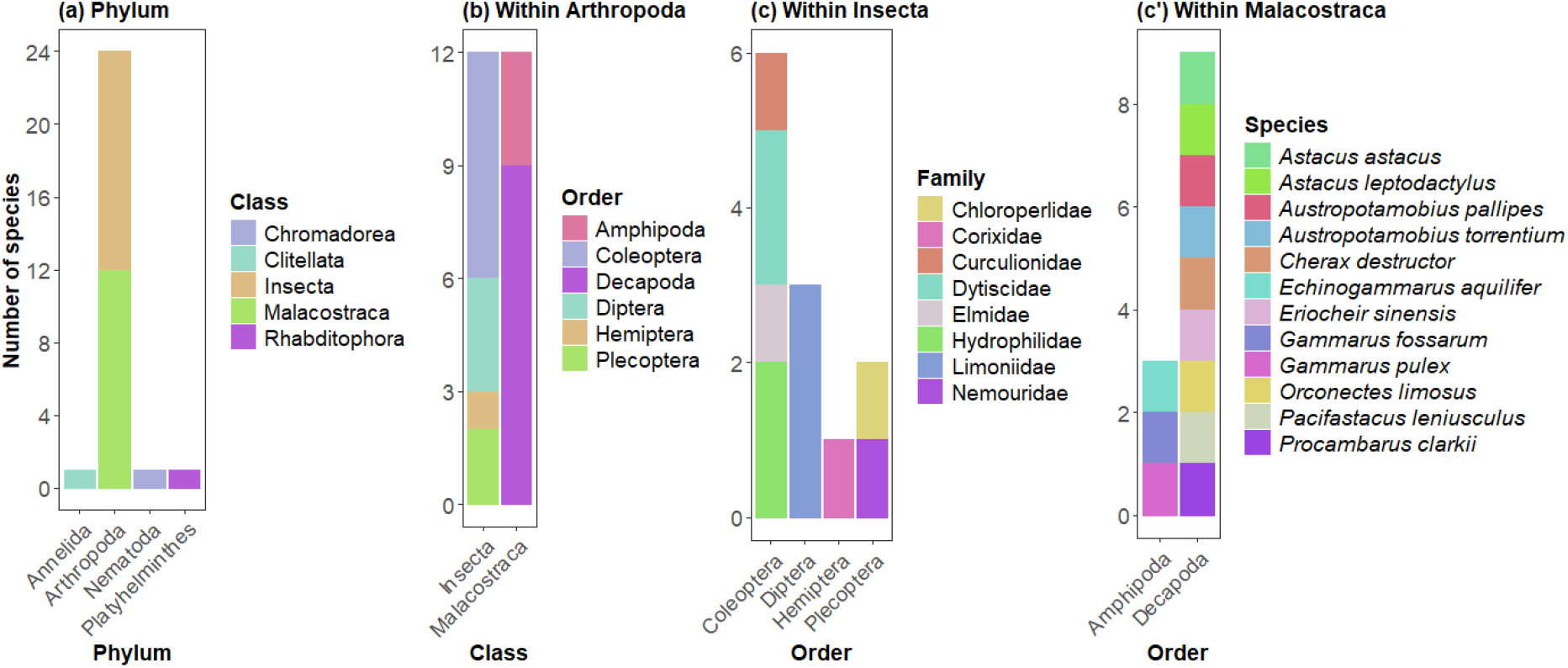
Gap analysis of the aligned DNA library. Number of species associated with COI-5P genetic sequences not alignable with fwh2 primer pair at the phylum rank (a), within the Arthropoda phylum (b) and within the Insecta and Malacostraca class (c, c’).

Arthropoda emerged as the predominant taxonomic groups facing alignment difficulties at the phylum level. Within insects (Arthropoda), Coleoptera and Diptera had the species with the most alignment issues. Specifically, within Malacostraca (Arthropoda), the gammarid (Gammaridae) and crayfish (Astacidea) families had the highest number of species with alignment issues (Figure 3). Several reasons could explain these results. Firstly, although COI-5P sequences were available, some were found to lie outside the primer pairs’ intended region during the alignment process, either entirely or partially. In the latter case, where the sequence in question was shorter than the desired amplified barcode, it was subsequently discarded. Another possible explanation is the poor sequence annotation, such as mislabelling (e.g. COI-3P instead of COI5-P) or misidentification of specimens, leading to incorrect species assignments. Misidentification of voucher specimens has been highlighted as a major factor contributing to erroneous records, as morphological identifications of closely related species can be challenging. This issue has been noted by Leite et al. 2020; Paz & Rinkevich, 2021; Pentinsaari et al. 2020, emphasizing its impact on subsequent species identifications using databases like BOLD. This was evident in cases when some species of insects failed to align with the primer pair, while others belonging to the same genus did. Despite BOLD being a curated database with verification procedures during sequence deposition and a reported error rate of less than 1% for Metazoan sequences at the genus level (Leray et al. 2019), problematic records may still exist, as evidenced in various studies on marine macroinvertebrates (Radulovici et al. 2021) where up to 39% of sequences were considered ambiguous. For the Decapoda species (Malacostraca, Arthropoda) that failed to align with the primers pair fwhF2/fwhR2N, comprising eight crayfish species and one crab species, their absence in the mock community during primer design could account for this discrepancy (Elbrecht and Leese 2015, Elbrecht et al. 2017). Although Gammarids (Amphipoda, Arthropoda) were included in primer design, the failure of some species to align with the primers could be due to their vast diversity within aquatic environments (Horton et al. 2023). Molecular studies on Amphipoda have revealed extensive species diversity and the presence of cryptic species complexes (Jażdżewska and Mamos 2019), suggesting a high genetic variability within species (Lefébure et al. 2006) that may not be compatible with the primer pair. Furthermore, Vamos et al. 2017 indicated that their analysis of the efficiency of the primers pair displayed higher penalty scores for certain taxa of Turbellaria, Mollusca, Trichoptera, and Isopoda, indicating potential underrepresentation due to primer bias. This aligns with findings from other studies suggesting preferential detection of taxonomic groups by different markers and primers (Leduc et al. 2019). Consequently, the incorporation of multiple genetic regions in DNA metabarcoding to ensure the broadest possible taxonomic detection may prove to be a good solution (Duarte et al. 2021). This could be particularly important to monitor the noble crayfish, *Astacus astacus* and the amphipod *Gammarus roeselii* which cannot be detected with fwhF2/fwhR2N primers pair, (Figure 3), whereas they are the most frequently monitored species of the malacostracans in Europe (Weigand et al. 2019). Analysis of the number of haplotypes relative to the number of species available within different phyla (Figure 4) provides important information about the potential to detect taxa in natural samples.

**Figure 4.**
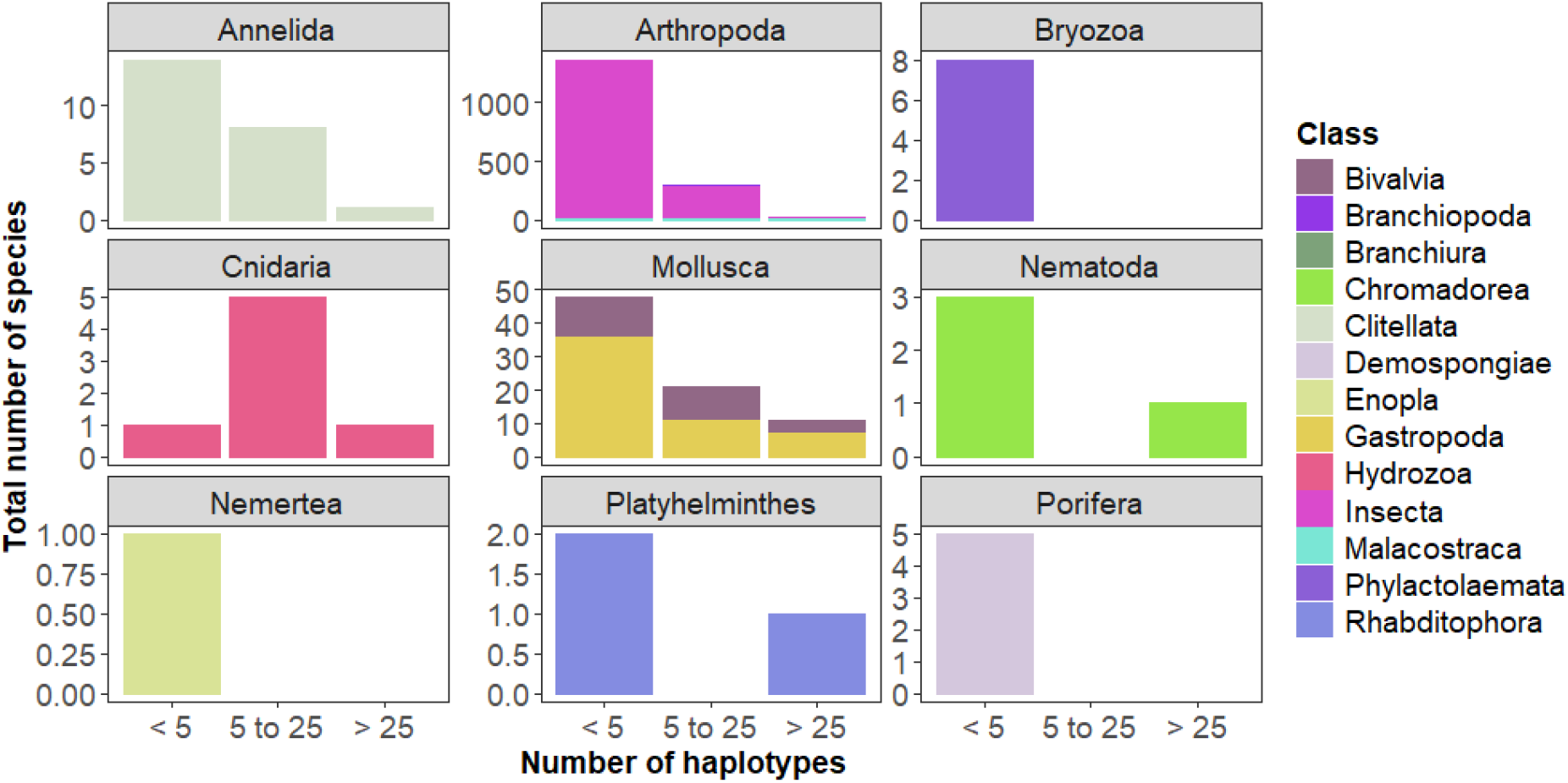
Taxonomic composition at the class rank for the species from the aligned DNA library. Each box at the phylum level represents the distribution of COI-5P haplotypes relative to the number of species.

In our reference library, for most phyla, the majority of species have less than five haplotypes, except cnidarians, which have the majority of their species with 5 to 25 haplotypes. Only three species presented more than 100 unique barcodes: one trichopteran (*Agraylea multipunctata*), one isopod (*Asellus aquaticus*) and one gasteropod (*Physella acuta*). These findings are consistent with the observations of Trebitz et al. 2015, suggesting a noteworthy exception to the prevailing low barcoding rate for invertebrates, with some species being exceptionally well genetically represented. This phenomenon may be attributed to the scientific significance and ecological relevance of these species, such as being acknowledged as a reliable indicator. Indeed, *Asellus aquaticus* is a common species monitored in European countries (Weigand et al. 2019), known as bioindicator for metal pollutants detection (e.g. O’Callaghan et al. 2019). *Physella acuta* is also intensively studied as it is an invasive aquatic Gasteropoda with worldwide distribution (Banha et al. 2014, Vinarski 2017). The analysis on the ability of the fwhF2/fwhR2N primers to discriminate each species by a unique sequence demonstrated that 57 identical sequences were shared by 116 distinct species (Table S2). Among those sequences, 10 were shared by two or three different genus and 47 were shared by two or three different species. These results could be explained by the fact that some taxa could have been mislabelled or that, for this length of barcode (205 pb), no genetic variability is found between two related taxa, therefore this barcode is not suitable to decipher those species. Several authors showed the necessity to have multiple sequences for each species to cover correctly their haplotypic diversity (Leite et al. 2020, Keck et al. 2023). This imperative arises from the recognition that the absence of intraspecific variants can pose significant challenges. In instances where a single sequence is available for a species exhibiting high genetic variability, the accurate identification of all haplotypes may be compromised. Moreover, inadequate representation of closely related species within reference libraries can lead to the erroneous assignment of multiple species to a single taxonomic entity (e.g. Jackman et al. 2021), potentially resulting in erroneous assessments of species diversity.

## Conclusions

Our study underscores a widespread absence of reference barcodes for numerous extant invertebrate species. Although barcoding offers advantages over morphological identification in biomonitoring, existing gaps in barcode libraries may hinder their effectiveness (Duarte et al. 2020, Feio et al. 2020, Hestetun et al. 2020, Vieira et al. 2021). Substantial efforts are necessary to sequence new individuals for species absent from in the reference barcoding library, and for species with a low number of sequences. This will enable a more efficient and reliable species identification and biodiversity assessment, since the effectiveness and reliability of DNA barcoding identification are linked to the thoroughness of taxonomic curation and completeness of the reference barcoding libraries (Geiger et al. 2021) (Keck et al. 2023). Nevertheless, our work has established a reference barcoding library for freshwater macroinvertebrates at the French level. This database can be utilized for biodiversity studies employing environmental DNA with short COI primers (fwhF2/fwhR2n) designed for samples containing partially degraded DNA. Furthermore, we have provided insights into the various biases associated with the utilization of this library, that future users will have to take into account to interpret their results. Looking ahead, an important future development for the fwhF2/fwhRNn reference library would be to assign a confidence level to the identification of each taxon (e.g. species). This confidence could be determined based on several factors, including the number of reference sequences available (with higher sequence counts providing greater confidence), geographical coverage (broader coverage being preferable), species delimitation (with monophyletic groups offering more robust identification compared to paraphyletic ones), and the availability of metadata associated with each sequence (e.g. sampling date and location, habitat, sequence quality). These criteria align with recommendations proposed by (Fontes et al. 2021).

## Acknowledgments

This study was made possible thanks to a funding of the Carnot Eau&Environnement institute, a funding of the INRAE department AQUA and a funding of the pôle INRAE-OFB ECLA (ECosystèmes LAcustres). We would also like to thank Tristan Lefébure and Maylïs Gauthier from LEHNA (CNRS) for their insights into this study. The authors have declared that no competing interests exist.

