## Supplementary figures and images for "An annotated reference library for supporting DNA metabarcoding analysis of aquatic macroinvertebrates in French freshwater environments"

### Figure S1

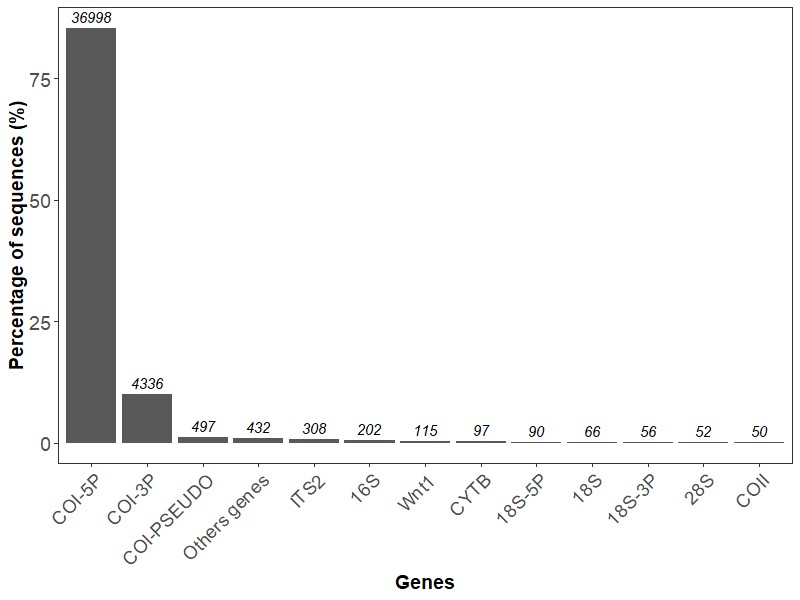
